# Polygenic scores for height in admixed populations

**DOI:** 10.1101/2020.04.08.030361

**Authors:** Bárbara D. Bitarello, Iain Mathieson

## Abstract

Polygenic risk scores (PRS) use the results of genome-wide association studies (GWAS) to predict quantitative phenotypes or disease risk at an individual level. This provides a potential route to the use of genetic data in personalized medical care. However, a major barrier to the use of PRS is that the majority of GWAS come from cohorts of European ancestry. The predictive power of PRS constructed from these studies is substantially lower in non-European ancestry cohorts, although the reasons for this are unclear. To address this question, we investigate the performance of PRS for height in cohorts with admixed African and European ancestry, allowing us to evaluate ancestry-related differences in PRS predictive accuracy while controlling for environment and cohort differences. We first show that that the predictive accuracy of height PRS increases linearly with European ancestry and is largely explained by European ancestry segments of the admixed genomes. We show that differences in allele frequencies, recombination rate, and marginal effect sizes across ancestries all contribute to the decrease in predictive power, but none of these effects explain the decrease on its own. Finally, we demonstrate that prediction for admixed individuals can be improved by using a linear combination of PRS that includes ancestry-specific effect sizes, although this approach is at present limited by the small size of non-European ancestry discovery cohorts.

## Introduction

Genome-wide association studies (GWAS) have proved remarkably successful at identifying the genomic basis of complex traits. For example, 3,290 genome-wide significant loci explain approximately 25% of the phenotypic variation in height in European ancestry individuals^1^. This highly polygenic architecture is a feature of most common diseases^2^. One approach to incorporate this information into clinical care is to use polygenic risk scores (PRS). PRS are simply sums of the risk alleles carried by an individual weighted by their effect sizes^3^. At least for some diseases (for example, coronary artery disease and breast cancer) PRS can identify individuals with clinically actionable levels of risk^4–7^.

One major limitation is that the majority of participants in GWAS used to derive PRS are of European ancestry^8,9^. Although many genome-wide significant GWAS hits do replicate in non-European ancestry cohorts^10–14^, overall the predictive power of PRS is lower and decreases with genetic distance from Europeans^15–18^. There are many possible explanations for this, including inter-cohort differences in data collection, phenotype or environment, differences in linkage disequilibrium (LD) structure or allele frequencies across populations^19^, differences in causal or marginal effect sizes, and epistatic or gene-environment interactions^20^. However, the relative contribution of these effects is unknown. Consequently, the clinical utility of PRS has largely been explored in European ancestry populations and little is known about how biological and methodological factors influence prediction in non-Europeans^4,15,21,22^. Moreover, while some studies have compared PRS performance across discrete ancestry groups, performance as a function of continuous ancestry – particularly important in recently admixed populations – remains largely unexplored.

Here, we address this gap in two ways. First, we describe the reduction in the predictive power of height PRS as a function of ancestry in populations of recently admixed African and European ancestry. Height is a well-studied model for understanding complex polygenic traits, and admixed populations allow us to investigate predictive power as ancestry varies continuously, while controlling for environmental or methodological differences between cohorts. Second, we explore the roles of different biological and statistical factors in driving this reduction. Our results suggest that there is no simple statistical solution to the problem of PRS transferability and emphasize the importance of performing GWAS in diverse populations.

## Subjects and Methods

### Data preparation, QC, and phasing

We obtained genotype and phenotype data from the UK Biobank^23^ (UKB), Women’s Health Initiative^24^ (WHI), Jackson Heart Study^25^ (JHS) and Health and Retirement Study^26^ (HRS) cohorts through dbGaP. For WHI and JHS we lifted over SNP positions to hg19 using *liftOver*. For WHI, JHS and HRS, we flipped alleles to the positive strand using the appropriate strand files from https://www.well.ox.ac.uk/~wrayner/strand/. We identified individuals with admixed African ancestry in each cohort using a combination of genetic clustering and self-reported ancestry as follows:

### UKB

This dataset contains several ancestry groups. We selected 8,813 individuals with African or admixed African and European ancestry based on PCA **(Figure S1)** and refer to them as UKB_afr. We further filtered this set to contain individuals with at least 5% of African ancestry, resulting in 8,700 individuals (**Table 1**). We randomly selected 9,998 European ancestry individuals from the “White British” subset to use as a comparison sample and refer to them as “UKB_eur”.

**Table 1.** Datasets used in this study. UKB, UK Biobank; WHI, Women’s Health Initiative; JHS, Jackson Heart Study; HRS, Health and Retirement Study; N, number of individuals; a, number of SNPs in the intersection between genotyped SNPs and SNPs from the height GWAS that passed our filters; b, SNPs clumped in 100 Kb windows and with p<0.0005; a, b, f, number of SNPs in imputed set in parentheses; d, randomly selected from the entire European component of the cohort. CI, approximate 95% confidence intervals obtained from bootstrapping.

| Dataset | Ancestry | $N^f$ | Total number of SNPs<sup>a</sup> | Number of SNPs in PRS<sup>b</sup> | Partial-$R^2$ (CI, %) | European Ancestry (%) |
| --- | --- | --- | --- | --- | --- | --- |
| UKB European subset (UKB_eur) | European | 9,998<sup>d</sup> | 685,475 | 6,052 | 22.4 (20.8-24) | 100 |
| HRS European (HRS_eur) | European American | 10,159 (10,123) | 1,515,431 (10,118,786) | 7,117 (9,724) | 15.6 (14.4-16.9) | 100 |
| UKB admixed African (UKB_afr) | African + European | 8,700 (8,696) | 685,475 (13,279,553) | 6,049 (10,577) | 4.1 (3.2-4.9) | 13.1 |
| WHI (WHI_afr) | African American | 6,863 | 741,983 | 5,744 | 3.6(2.8-4.5) | 22.7 |
| JHS (JHS_afr) | African American | 1,773 | 702,685 | 5,676 | 3.8 (2.2-5.7) | 17.7 |
| HRS admixed African (HRS_afr) | African American | 2,251 (2,241) | 1,511,742 (10,118,786) | 7,101 (9,724) | 3.1 (1.9-4.6) | 17.5 |

### WHI

This dataset contains both African American and Hispanic participants. We ran unsupervised ADMIXTURE^27^ with k=3 and identified 7,285 individuals with self-reported “African American” ancestry with at most 0.8 of the first ADMIXTURE component (which we interpret as reflecting European ancestry), and at most 0.05 of the second (which we interpret as reflecting Native American ancestry; **Figure S2)**. We further filtered this set to contain individuals with at least 5% of African ancestry and height between ±2 standard deviations (sd) from the mean (see **Figure S12**), resulting in 6,863 individuals (**Table 1**). We refer to them as “WHI_afr”.

### HRS

This dataset contains multiple ancestry groups, but oversamples African American and Hispanic participants. We ran unsupervised ADMIXTURE^27^ with k=3, and identified 2,322 individuals with self-reported “Black/African American” ancestry with at least 0.05 of the first ADMIXTURE component (interpreted as reflecting European ancestry), at most 0.05 of the second ADMIXTURE component (interpreted as reflecting Native American ancestry, see boxed area in **Figure S3).** We further filtered this set to contain individuals with at least 5% African ancestry and height not less than 2 sd below the mean (for each sex, **Figure S12**), resulting in 2,270 individuals (referred to as “HRS_afr”). We also identified 10,486 individuals who self-reported “White/Caucasian” ancestry and had at most 0.05 of each of the first two ADMIXTURE components (“HRS_eur”), of which 10,159 had sex-specific height above the −2 sd cutoff (**Figure S12, Table 1**).

### JHS

This dataset contains only African American participants, so we did not filter the data further based on genetic clustering. We kept all 1,773 individuals with at least 5% African ancestry (“JHS_afr”).

### GWAS results

We obtained UK Biobank summary statistics for height from the Neale Lab GWAS (round 2; https://www.nealelab.is/uk-biobank). We used a set of 13,586,591 autosomal SNPs that passed QC filters of INFO score > 0.8 and MAF > 0.0001 For some analyses we used between-sibling effect sizes estimated at a subset of 1,284,881 SNPs^28^. The number of individuals and SNPs per dataset are shown in **Table 1**.

### Ancestry inference

For the admixed cohorts, we estimated local ancestry for each individual and computed the genome-wide proportion of European ancestry. To do this, we merged each dataset with CEU (Utah residents with Northern and Western European ancestry) and YRI (Yoruba from Ibadan, Nigeria) individuals from the 1000 Genomes Project^29^ and phased each dataset separately using HAPI-UR^30^ with a window size of 91. We then used RFMix^31^ to infer local ancestry, using the CEU and YRI individuals as references for European and African ancestry, respectively. We used the most likely ancestry path inferred by the Viterbi algorithm of RFMix to estimate proportions and checked that they were consistent with those obtained from unsupervised ADMIXTURE with k=2 **(Figure S4)**.

### Clumping and pruning SNPs

We first intersected the ∼13.5 million SNPs with UK Biobank summary statistics and the genotyped SNPs in each dataset **(Table 1)**. Next, we defined sets of SNPs based on a variety of clumping strategies. We clumped SNPs in physical and genetic windows using a range of p-value thresholds. Physical window sizes (in Kb) were: 1,000, 500, 100, 75, 50, 25, 10, 5. Genetic window sizes (in cM) were: 1, 0.5, 0.3, 0.25, 0.2, 0.15, 0.1. We considered SNPs below the p-value thresholds: 5×10^−7^, 5×10^−6^; 5×10^−5^ 5×10^−4^, 5×10^−3^. For each set of parameters, we followed these steps: 1) retain only SNPs below the p-value threshold, 2) select lowest p-value SNP, 3) remove SNPs within the window around that SNP, 4) repeat steps 2 and 3 until there are no SNPs left. We also used a strategy of clumping based on empirical LD structure. We used PLINK2^32^ to estimate *r*^2^ between pairs of SNPs using UKB_eur (--clump-p1 0.01 –clump-r2 0.5 --clump-kb 250| 100| 50). Finally, we also applied a strategy where we clumped SNPs in approximately independent LD blocks^33^ (defined in either African or European populations). In total, we evaluated 80 pruning strategies **(Table S1)**.

For the unweighted PRS, we tested prediction for the same 80 sets of SNPs **(Table S1)**. For analyses using effect sizes re-estimated from sibling pairs and imputed genotypes, we repeated these steps, except restricting the initial set of SNPs (before pruning/clumping) to those present in the sibling or imputed dataset. For imputed genotypes, we performed the 40 clumping strategies using physical windows described above **(Table S1)**.

### Effect size estimates for African ancestry

We ran a GWAS using PLINK^34^ on the Admixed African individuals from the UK Biobank including sex, age, age^2^, and the first 10 principal components, computed using smartpca^35^, as co-variates. We then computed a chi-squared statistic for the difference between the Admixed African effect size (*β* _*afr*_, with standard error *σ*_*afr*_) we obtained and the European effect sizes from the UK Biobank (*β*_*eur*_ with standard error *σ*_*eur*_):

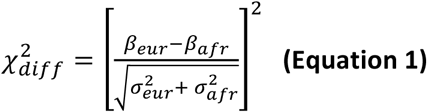

### PRS and odds-ratio calculation

We calculated PRS for each individual, *j*, as the weighted sum of effect sizes:

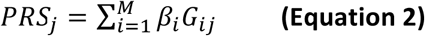

where the sum is over all *M* SNPs used in the PRS, *G*_*ij*_ is the effect allele dosage (0, 1 or 2) of individual *j* at SNP *i*, and *β*_*i*_ is the estimated effect size of the effect allele at SNP *i*. To calculate unweighted PRS, we set *β*_*i*_ to ±1 depending on whether the original *β*_*i*_ is positive or negative.

To evaluate predictive power, we fitted a linear model of height as a function of sex, age, age^2^, genome-wide European ancestry (*p*_*eur*_) and PRS (*height*∼*sex + age^2^ + p*_*eur*_ *+ PRS*), and compared it to a model without PRS (*height*∼*sex + age + age^2^ + p*_*eur*_). The partial-*R*^*2*^ between the two models gives the proportion of the phenotypic variation explained by the PRS, which we refer as partial-*R*^2^ or predictive power, throughout. We estimated standard errors using 1,000 bootstrap replicates with replacement. To evaluate the effect of ancestry on predictive power, we stratified each dataset into 2-4 equally sized bins based on the proportion of European ancestry of individuals. Next, we calculated the partial-*R*^2^ for each bin and standard errors estimated from 1,000 bootstrap replicates. Finally, we performed a weighted regression–using the inverse of the bootstrap variance as weights—of the partial-*R*^2^ values on the median proportion of European ancestry in each bin. We repeated this analysis with imputed genotypes, unweighted PRS and sibling-estimated effect sizes.

We also constructed linear combinations of *PRS*^17^. Using **Equation 2**, we calculated *PRS*_*eur*_ using effect sizes from the UK Biobank, and *PRS*_*afr*_ using the same SNPs as *PRS*_*eur*_ but with effect sizes we re-estimated from the UKB_afr dataset (see above). In 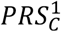, we weight *PRS*_*afr*_ in all individuals by a common factor α ranging from 0-1, and in 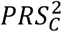, in addition to α, each individual’s *PRS*_*afr*_ is weighted by *p*_*eur*_, the proportion of European ancestry for the individual. So, for each individual *j*:

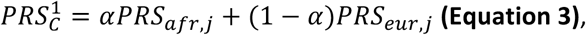

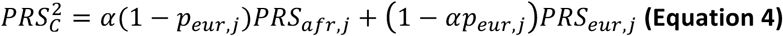

We evaluated the predictive power of 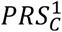 and 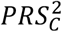 in WHI_afr, JHS_afr and HRS_afr **(Figure 2)**.

**Figure 1.**
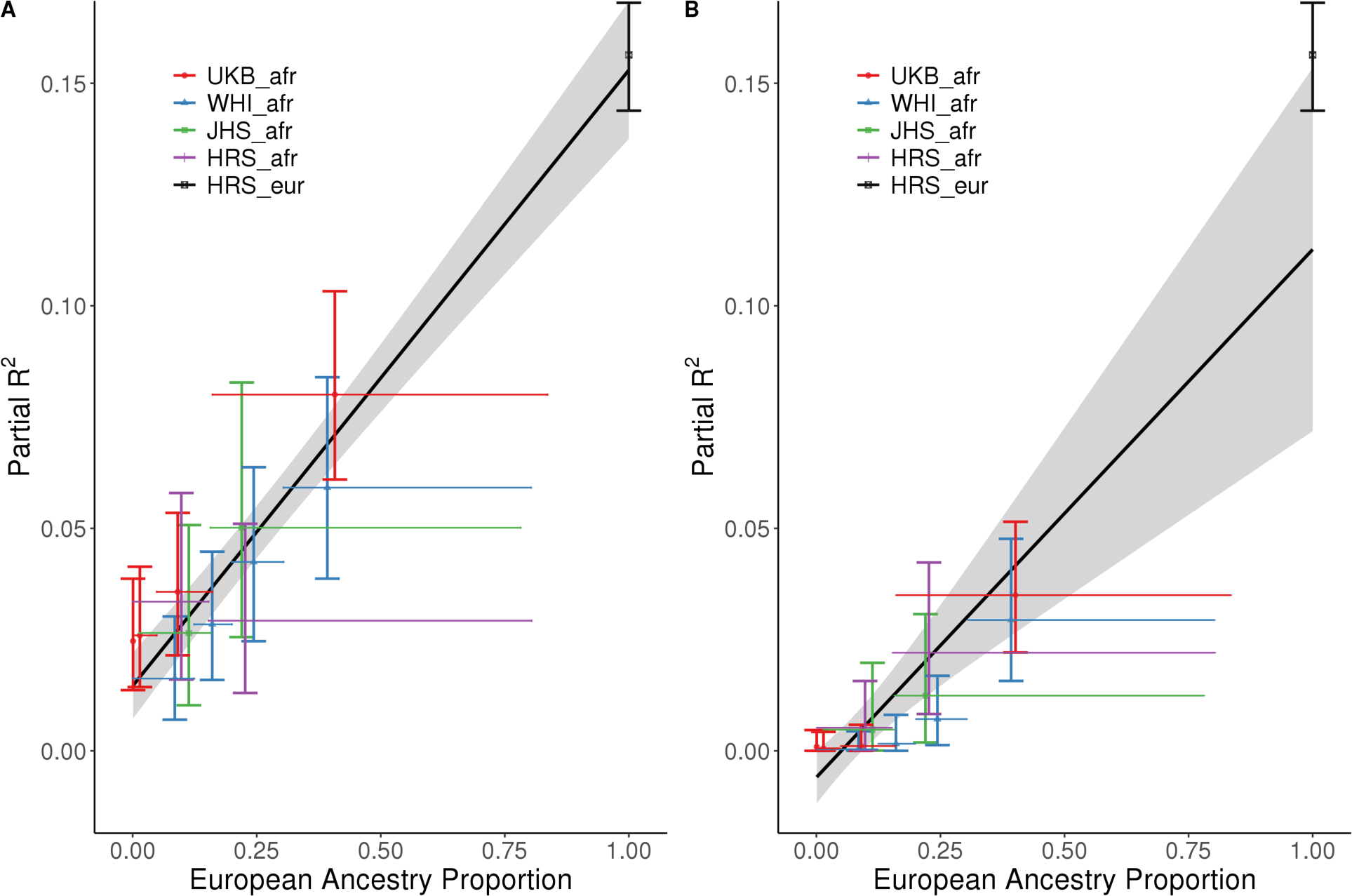
Partial-*R*^2^ as a function of European ancestry in admixed populations. Each admixed dataset is split up into quantiles of ancestry. Each cross represents a single quantile bin. Vertical bars represent bootstrap confidence intervals. Horizontal bars represent the range of European ancestry proportion included in bin. **A**: Using all segments of the genome; **B**: using only European ancestry segments.

**Figure 2.**
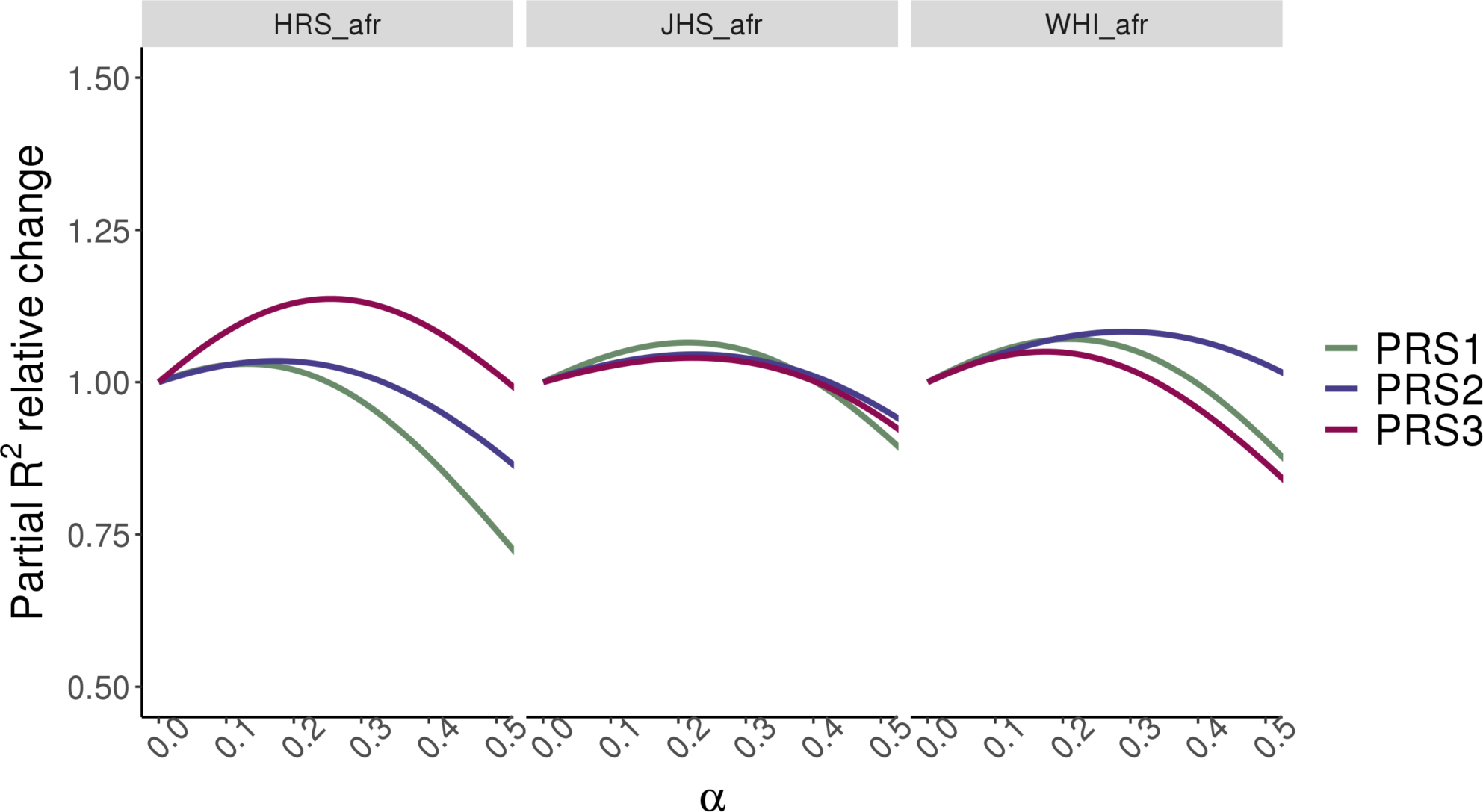
Predictive power of linear combinations of PRS. Relative increase in partial-R^2^ for HRS_afr (N=), JHS_afr (N=), WHI_afr(N=) from three linear combination of PRS_eur_ and PRS_afr_. For 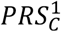 and 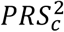, α represents the constant weight given to the African component across individuals. 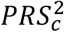, in addition to α, weights the African component based on individual African ancestry 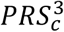 uses European effect sizes for PRS effect alleles falling in European ancestry segments, and a linear combination of European and African betas (weighted by α) for PRS effect alleles falling in African ancestry segments (see Equations 3-5).

We also used **Equation 2** to calculate PRS based only on European ancestry segments of the genome and repeated the analysis of partial-*R*^*2*^ as a function of European ancestry.

Finally, we constructed a combined PRS where Admixed African effect sizes are used for SNPs falling in African ancestry regions of the genome, and European effect sizes are used for SNPs falling in European ancestry regions. For African ancestry segments, effect sizes from admixed Africans are weighted by a constant, α. So, for each haplotype in each individual, we have:

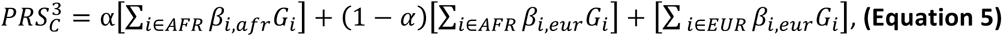

where *G*_*i*_ is the genotype of the *i*-th SNP, and *EUR* and *AFR* are the sets of European and African ancestry regions of the genome. We then sum 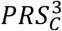 for both haplotypes, and refer to this sum as 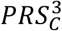 for simplicity.

We estimated the odds ratio for being in the upper *q*-quantile of height conditional on being in the upper *q*-quantile of PRS as:

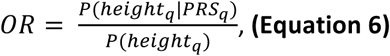

where *P*(*height*_*q*_) is the proportion of individuals in the upper *q*-quantile of height and *P*(*height*_*q*_|*PRS*_*q*_) is the proportion of individuals in the upper *q*-quantile of PRS who are also in the upper *q*-quantile of height. We used standardized residuals of height after regressing out *age, age*^*2*^, *sex* for each dataset.

### Recombination rate and LD score analyses

We computed genetic distance using recombination maps estimated in African Americans (AA_Map)^19^ using linear interpolation between genotyped points to estimate genetic distance in 20 Kb windows. We stratified each dataset into four equally sized bins according to recombination rate distance and calculated partial-*R*^2^ and bootstrap confidence intervals (3,000 replicates) for each bin. We then divided the values for each bin by those obtained for the full dataset, thus obtaining a relative partial-*R*^2^. In another approach we tested for correlation between these genetic distances and 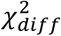 **(Equation 1)** and LD scores^36^.

### Genetic and phenotypic variance analyses

We estimated the ratio of the additive genetic variance explained by the PRS SNPs as:

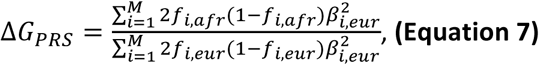

where *f*_*i,eur*_, *f*_*i,afr*_, *β*_*i,eur*_ and *β*_*i,afr*_ are the allele frequencies and effect sizes for each of the *M* PRS SNPs in the European or admixed African ancestry cohorts, respectively. For HRS_afr, HRS_eur, UKB_afr and UKB_eur, allele frequencies were obtained directly from the datasets. For WHI_afr and JHS_afr, we compared to “European” frequencies from the CEU individuals from Phase 3 of the 1000 Genomes Project^29^.

### Modeling height variance as a function of ancestry

We combined all 29,746 individuals **(Table 1)** and computed the residuals *y*_*i*_ of the regression of height on *sex, dataset, age, age*^*2*^, *sex*dataset, sex*age, sex*age*^*2*^, *dataset*age, dataset*age*^*2*^. We then fitted a linear model for residual height as a function of the ancestry of individual *j* (*p* _*i, eur*_) but also allowing the variance to linearly with ancestry:

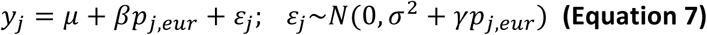

for model coefficients *μ, β, σ*^2^ and *γ*. We fit this model using the GAMLSS package^37^ in R.

### Local differences in allele frequency

We calculated allele frequencies for all variants in the HRS_afr and HRS_eur subsets separately. We defined 6 and 10 Kb windows around each PRS SNP and calculated the mean squared frequency difference between both subsets for all the SNPs contained in the window. We explore the effect size difference for AFR and EUR **(Equation 1)** for each PRS SNP as a function of the mean squared frequency difference in the window.

## Results

### Constructing polygenic risk scores

We tested 80 approaches to PRS construction. Broadly, we observe that including more SNPs in the PRS increases partial-*R*^2^ (rho=0.36 for UKB_eur, p= 0.0011), although this masks subtler patterns **(Figure S5 and Table S1)**. For example, when using a clumping strategy with a fixed window size, increasing the p-value threshold (more SNPs) leads to higher predictive power. On the other hand, when using a fixed p-value, increasing window sizes (fewer SNPs) also improves prediction **(Figure S5)**. This is consistent with previous findings that avoiding redundant information from non-causal SNPs can be more important than including all causal SNPs^16,38,39^. The LD clumping strategies^5,16,22^ have the highest predictive power (e.g. 18.9% in HRS_eur for a 250 kb window, Table S1), but also includes 1.4 times as many SNPs as any other strategy we tested **(Table S1)**. The approximately independent LD block strategies^33^ yield small sets of SNPs that explain almost as much variation as the much (23-37 times) larger LD clumping sets, but also rely on prior knowledge about the population-specific LD structure. The LD-based pruning and clumping strategies also show the highest difference in PRS between 1000 Genomes European and African Populations **(Table S1)**.

Thus, we focused on strategies that are independent of LD and chose a set of SNPs based on a p-value threshold of 0.0005 and a physical window of 100 Kb, which includes ∼5,600-7,100 SNPs **(Table 1)** obtains partial-*R*^*2*^ values close to the LD clumping strategies while requiring about 10-fold fewer SNPs **(Table S1, Figure S5).** Nonetheless, results for other strategies are qualitatively similar **(Table S1)**.

### Predictive power increases linearly with proportion of European ancestry

We estimated the predictive power of the PRS in each dataset. Partial-*R*^*2*^ was 22.4% in UKB_eur, and 15.6% in HRS_eur **(Table 1)**. Because the 9,998 UKB_eur individuals analyzed here were also in the discovery GWAS, we use the HRS_eur dataset throughout the paper as representative of European ancestry. In the admixed cohorts (WHI_afr, JHS_afr, HRS_afr, UKB_afr), partial-*R*^*2*^ was 3.1%-4.1%, or between 3.8 to 5-fold lower than in HRS_eur, consistent with previous observations^15,16^.

Stratifying individuals in each cohort by their average genome-wide ancestry, we find that partial-*R*^2^ increases linearly with European ancestry (by 1.3% for each 10% increase in European ancestry; **Figure 1A**). We estimated the partial-*R*^2^ in individuals with no European ancestry (i.e, the intercept of this regression) to be 1.5% (S.E.=0.3%). This result is robust to the set of SNPs used in the PRS, with intercepts ranging between −1% and 2.5%, depending on pruning strategy **(Figure S5)**. Relevant for clinical interpretation, the odds-ratio for ‘tallness’ in the tails of the PRS distribution is also lower in the admixed populations than in the European ancestry population, although only 2.3-fold on average across populations between the highest and lowest 5% of the European ancestry spectrum (95^th^ quantile of PRS distribution), compared to the 3.8 to 5-fold difference in partial-*R*^*2*^ **(Figures S7, S8)**. We also see this linear trend when we restrict the PRS SNPs to those found in regions of the genome inferred to have European ancestry **(Figure 1B)**. Consistent with previous results ^40^, we conclude that the predictive power of the PRS comes largely, but not entirely, from the European ancestry segments of the admixed genomes.

Next, we explored whether combining ancestry-specific PRS could improve predictive power, as suggested by Ref. 17. A simple linear combination of PRS improves partial-*R*^2^ in WHI_afr from 3.6% to 3.9%, in JHS_afr, from 3.8% to 4.1% and in HRS_afr from 3.1% to 3.2% for **(Figure 2)**. Weighting the combination by the ancestry proportion of each individual produces a similar improvement: 3.9% for WHI_afr, 4% for JHS_afr and 3.2% for HRS_afr **(Figure 2)**. Finally, we used local ancestry information to construct a PRS using ancestry-specific effect sizes at each SNP **(Methods, Figure 2)** This produces a similar improvement to the global ancestry weighted PRS, with partial-*R*^2^ increase between 0.1 and 0.3% across datasets. While these absolute improvements are modest this is likely due to the discrepancy in the GWAS sample size (N=8,813 Admixed African and N=361,194 European). With larger African ancestry GWAS, we expect that we would be able to improve the PRS performance in the admixed populations with this approach.

### Why does predictive power vary with ancestry?

A number of explanations have been proposed to explain why the predictive power of PRS is lower in non-European ancestry populations. These include differences in LD patterns, allele frequency of PRS variants, additive genetic variance, gene-gene (G×G) and gene-environment (G×E) interactions in different populations, as well as effect size estimation (for example, biases in the discovery cohort, differences in marginal effect sizes across populations or different causal variants). In this section, we evaluate the impact of some of these factors on PRS-based phenotypic prediction.

### Population-specific linkage disequilibrium (LD) patterns

Variants identified by GWAS are not generally themselves causal but rather are linked to the causal variant(s). Linkage disequilibrium (LD) patterns are known to differ between populations, suggesting that tag SNPs discovered in the original European ancestry GWAS may be less effective at tagging the causal variants on non-European ancestry haplotypes. This would result in reduced partial-*R*^2^ in admixed African populations when compared to European ancestry populations.

If LD differences between African and European haplotypes drive the pattern seen in **Figure 1** then a PRS constructed from SNPs in low recombination regions should be more transferable than a PRS constructed from SNPs in high recombination regions of the genome. When we bin PRS SNPs into quartiles of recombination distance and calculate PRS for SNPs in each bin, we see that partial-*R*^2^ for the admixed cohorts tends to decrease between the first and 4^th^ quantiles of recombination **(Figure 3B)**, suggesting that differences in LD do play a role in reducing prediction in non-Europeans. On the other hand, we note that, even for the quartile of lowest recombination, the reduction in partial-*R*^2^ is substantial–on average 76% across datasets–compared to 84% for the fourth quantile **(Figure 3A)**. Thus, even if all PRS variants were from low recombination regions, we would still observe a substantial reduction in predictive power.

**Figure 3.**
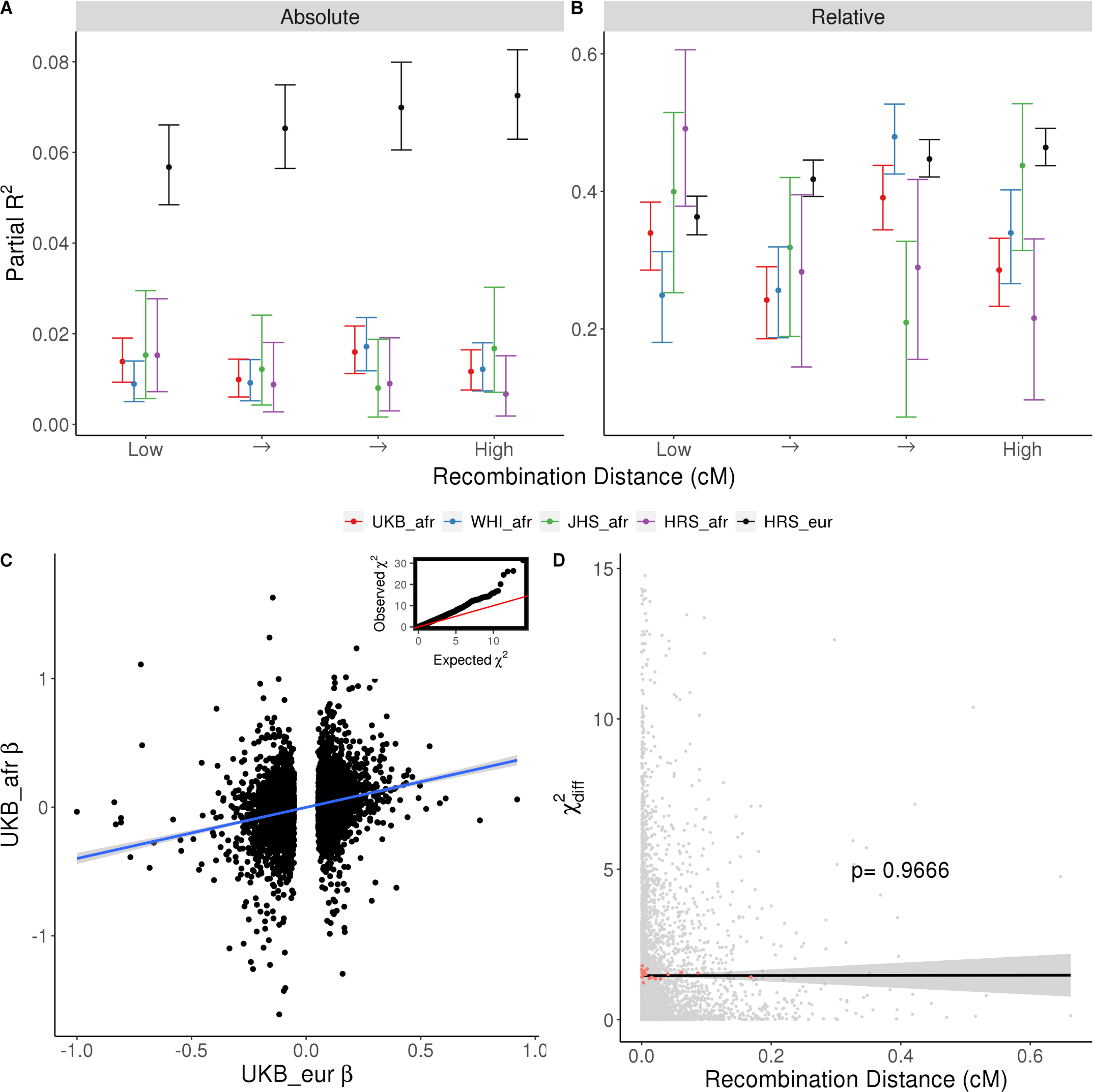
Effect of recombination rates on predictive power. **A** and **B**: PRS SNPs were binned into quantiles of PRS SNPs recombination distance in centimorgans (cM) according to an African Americans recombination map. y-axis, partial-R^2^, absolute (**A**) and relative (**B**), obtained for subsets of SNPs divided by the total partial-R^2^ for each dataset (**Table 1**). Confidence intervals were obtained by bootstrapping and dividing by total confidence intervals. **C**: correlation between PRS SNPs effect sizes from Europeans and Admixed Africans. Inset shows a qq-plot of 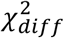 for PRS SNPs. **D**: y-axis, 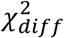 statistic for the difference in β between European and Admixed African populations (**Equation 1**), x-axis, recombination distance in cM. Cut-off at 15 for display purposes excludes 10 data points.

A second prediction is that the difference between effect sizes estimated in European and African ancestry populations should be larger in regions of high recombination. To test this, we evaluated whether effect sizes estimated directly from admixed individuals are different from the original (European) effect sizes **(Figure 3C)** and whether these differences are correlated with recombination in the regions in which they are located. We find no significant correlation (p=0.97, ρ=0.0005) between 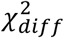 and local recombination rate **(Figure 3D)**, and a small positive correlation between 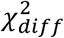 and European LD scores^36^ (p=0.0379,rho= 0.0292) **(Figure S10)**.

A further prediction is that, if differences in partial-*R*^2^ are driven by differences in ability to tag the causal variant, then PRS constructed from imputed genotypes should see less decrease in predictive power than those constructed from genotype array data. Using imputed genotypes for the HRS cohort, we find that the relationship between ancestry and partial-*R*^2^ is the same for imputed and array data suggesting that this is not the case **(Figure 4A)**. In fact, the absolute performance of imputed and array data is similar **(Figure 4B)**, consistent with previous observations^16^. This suggests that the genotyping array–in the HRS dataset—is efficient at capturing the SNP heritability. It is important to note that different datasets use different arrays and a different pattern could be observed for those datasets. We conclude that, while differences in LD do affect PRS transferability, they are not the only factor affecting relationship between ancestry and predictive power.

**Figure 4.**
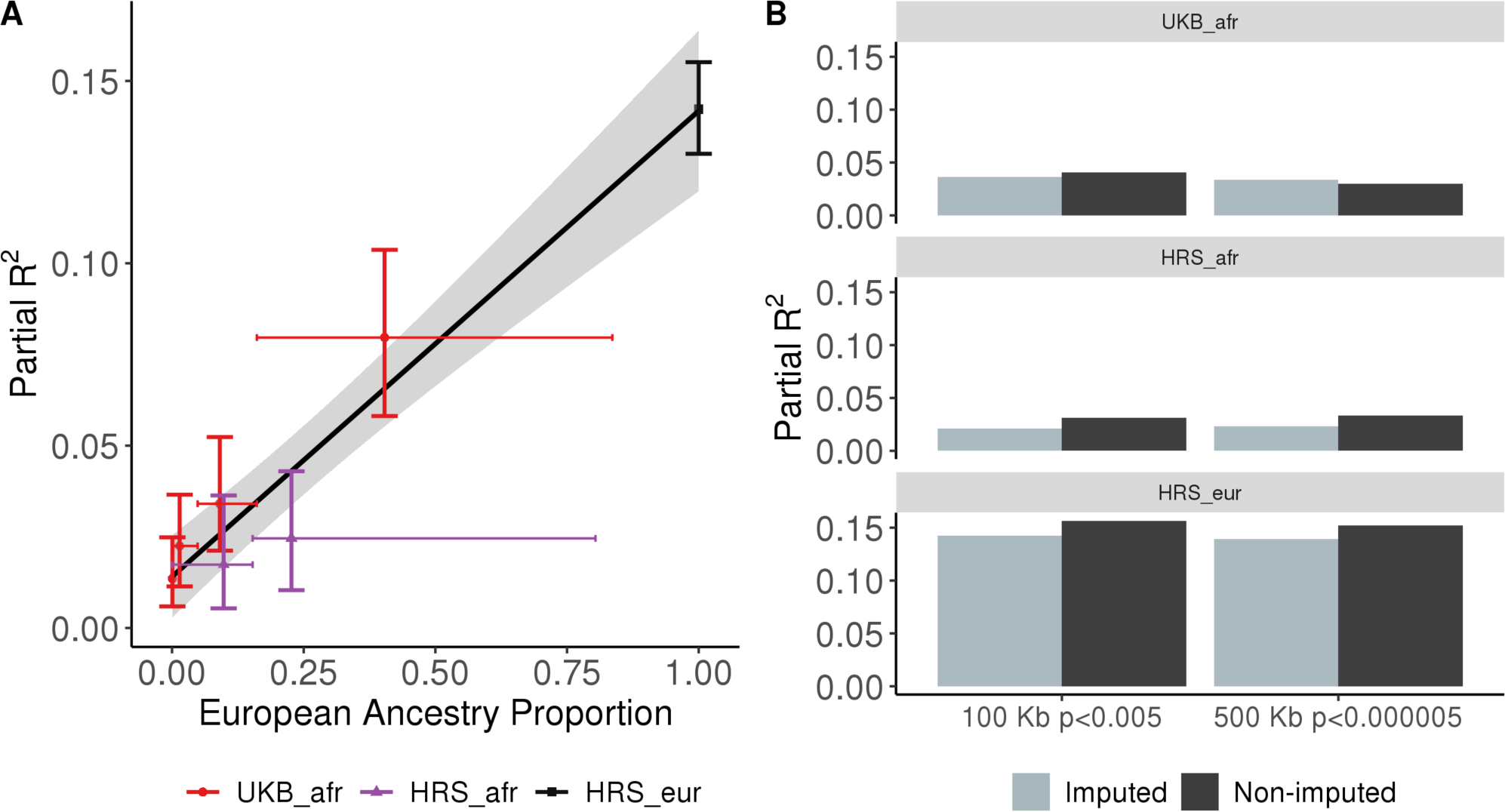
Imputed data. **A**: Partial-R^2^ as a function of European ancestry, where each admixed dataset is split up by quantiles of ancestry. As in **Figure 1**, each cross represents a single quantile bin. Vertical bars represent bootstrap confidence intervals. Horizontal bars represent the range of European ancestry proportion included in that bin. **B:** Partial-R^2^ for two clumping strategies (100 and 500Kb windows with p<0.005 and p<0.00005, respectively for imputed and genotyped sets of SNPs.

### Residual population stratification in discovery cohort

Despite statistical methods to control for population stratification, it continues to be a confounding factor in the analysis of GWAS results^22,41,42^ and could inflate predictive power in European relative to non-European ancestry cohorts. To test this, we used effect sizes at PRS SNPs re-estimated within sibling pairs from the UK Biobank^28^. This approach should remove much of the effect of population. We find that the linear relationship between ancestry and predictive power is similar to that observed for the GWAS PRS (**Figure S6**), albeit absolute partial-*R*^2^ values are lower across all datasets **(Figure S6, Table S1)**. We conclude that residual population structure in the UK Biobank GWAS results does not drive differences in predictive power across ancestries.

### Differences in the genetic variance captured by PRS SNPs

Differences in the frequencies of the tag variants could lead to different partial-*R*^2^ values across ancestries. Because GWAS have more power to detect more common variants, one hypothesis is that the PRS will tend to contain variants that are more common in European than African ancestry backgrounds–resulting in systematically lower predictive power in African ancestry populations. We tested this by comparing the ratio of the additive genetic variance contributed by the variants used in the PRS in the European and the admixed datasets **(Equation 7)**. We estimate this ratio to be 0.78 for UKB and 0.92-1.07 across the other datasets **(Table 2)**, suggesting that at most 8% of the decrease in partial-*R*^*2*^ (in non-UKB samples) can be explained by differences in the SFS. It is important to note that for the genotype data for JHS and WHI are less dense than for HRS and UKB **(Table 1**). One possibility is that those arrays are more biased towards SNPs that are common across ancestries.

**Table 2.** Expected reduction in predictive power due to differences in allele frequency spectrum. G_PRS_ ratio for PRS SNPs **(Methods)**. It is obtained by dividing G_PRS_ for the dataset-specific sum over all SNPs of 2p (1 − *p*)*β*^2^ in the admixed subset by the European subset. For WHI and JHS, allele frequencies from the CEU population from the 1000 Genomes were used as a proxy for the European component.

| Dataset | $G_{PRS}$ ratio |
| --- | --- |
| UK Biobank | 0.78 |
| WHI | 1.07 |
| JHS | 1.04 |
| HRS | 0.92 |

### Differences in the total genetic variance

A related possibility is that differences in site frequency spectrum might affect not just the variance explained by the PRS SNPs, but also the total genetic variance. Genome-wide heterozygosity in European ancestry populations is 30% lower than in West African ancestry populations^43^. If this were true for SNPs that causally affect height, then the additive genetic variance of those SNPs would also be 30% lower. Assuming constant environmental variance, it would follow that that the European phenotypic variance would be about 24% lower (0.8*0.7+0.2, assuming a heritability of 80%). Furthermore, the phenotypic variance in admixed populations would vary linearly with ancestry. In this case, the PRS could capture the same absolute amount of phenotypic variance, but the proportion of variance explained would be higher in European ancestry populations. However, we find that phenotypic variance does not vary significantly with genome-wide ancestry proportion once we regress out sex, age, age^2^, dataset and all their interactions (p=0.133, **Figure S11**). Therefore, differences in phenotypic and genetic variance cannot explain the pattern observed in Figure 1.

### Differences in marginal effect size

The marginal effect size at a PRS SNP depends on the cumulative effects of the causal variants that it tags. Therefore, marginal effect sizes at PRS variants across ancestries might differ for a number of reasons, including local epistasis or allelic heterogeneity. When we ignore effect sizes and calculate the unweighted PRS we see a very similar pattern to Figure 1 (**Figure 5A)**. This suggests that not only the marginal effect sizes, but even effect directions may differ between ancestries. That we can improve the predictive power of PRS by including effect sizes re-estimated in African ancestry populations **(Figure 2)** also indirectly supports the role of effect size differences. Finally, we find that allele frequencies tend to differ more between African and European populations around SNPs with larger effect size differences, although the effect is rather small (p=0.0327; rho= 0.0005; **Figure 5B).** This is observed for a larger window around the PRS SNPs as well **(Figure S9).** Taken together these results suggest that marginal effect sizes differ across ancestries, and that this is one of the factors underlying the reduction in predictive power.

**Figure 5.**
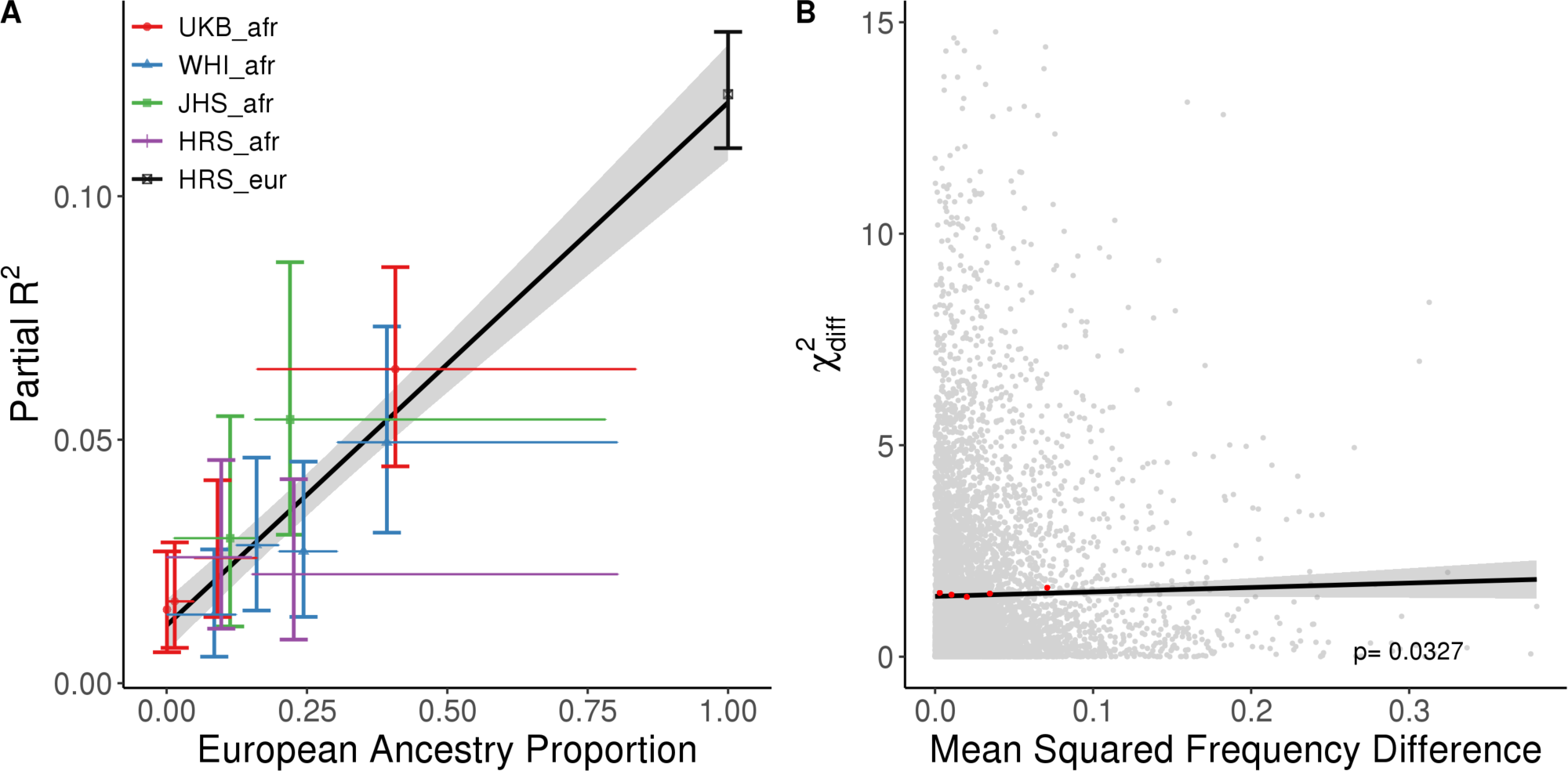
Unweighted PRS and the effect of local allele frequency differences on effect size differences. **A**: Partial-R^2^ for an unweighted PRS that uses the sign but not the magnitude of each SNP effect (**Methods**). Each admixed population is split up by quantiles of ancestry. Each cross represents a single quantile bin. Vertical bars represent bootstrap confidence intervals. Horizontal bars represent the range of European ancestry included in bin. **B**: X-axis, mean squared frequency difference for PRS SNPs for EUR and AFR in a 6 Kb window around each PRS SNP. Frequencies were calculated per dataset (HRS_eur, HRS_afr) for the causal allele. Y-axis, 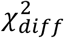 statistic for the difference in betas between EUR and AFR. Cut-off at 15 for display purposes excludes 15 data points. Black line is the linear regression with confidence intervals in gray. In red, for visualization, are median squared frequency difference values for 5 bins of mean squared difference and the mean χ^2^ for each bin.

## Discussion

Polygenic scores may become an important tool in translational and precision medicine, but are limited by their lack of applicability in non-European ancestry populations^9,22^. As a consequence, much of the potential of genomic disease risk profiling is restricted to European ancestry populations. Here, we show that the predictive power of PRS is approximately proportional to ancestry in populations of admixed European and African ancestry. We show that differences in LD structure and SFS do affect the transferability of PRS but do not, by themselves, explain the magnitude of the decrease.

Our results are broadly consistent with a recent estimate that LD and allele frequencies together explain up to 70% of the loss of accuracy in prediction between Europeans and Africans^44^, as well as empirical estimates that the trans-ancestry correlation in effect sizes for height is less than 1 (∼71-78%)^44,45^ and therefore that the marginal effect sizes at PRS SNPs are systematically different across ancestries. We interpret this as evidence that *cis*-epistasis or allelic heterogeneity–which mimics epistasis^46^–contribute to these differences. However, this may not, in general, be the only contributing factor. Gene-by-environment^45,47^ and gene-by-ancestry interactions may also contribute, and the relative importance of these mechanisms remains to be quantified.

By incorporating effect sizes from admixed populations in a linear combination of PRS, we are able to improve predictive power, in agreement with Ref. 17. Although inclusion of individual and local ancestry information yielded only a modest increase in prediction power, this is almost certainly due to our low sample size relative to number of markers^45^. With better-powered GWAS to estimate ancestry-specific effect sizes, the improvement should be more extensive. However, this suggests that large cohorts of diverse ancestries are needed in order to make PRS applicable to diverse ancestry groups and admixed populations. Leveraging information about the local ancestry background of each associated variant is a promising way to improve transferability, albeit that, too, requires larger non-European cohorts to estimate effect sizes. Finally, though we focused on cohorts of recent admixed European and African ancestry, additional work is required to characterize the transferability of PRS both in populations with more complex recent admixture, and in populations that are more anciently admixed.

## Supplemental Data

Supplemental data include 12 figures and one spreadsheet containing 20 tables.

**Table S1-1:6.** Different SNP sets used for weighted PRS calculation.

**Table S1-6:12.** Different SNP sets used for PRS calculation using effect sizes estimated from sibling pairs.

**Table S1-13.** Different SNP sets used for PRS calculation using imputed data.

**Table S1-14:19.** Different SNP sets used for unweighted PRS calculation.

**Table S1-20.** Difference between PRS values for 1000G super-populations for different sets of SNPs.

## Acknowledgments

This research was supported by a Research Fellowship from the Alfred P. Sloan foundation [FG-2018-10647], a New Investigator Research Grant from the Charles E. Kaufman Foundation [KA2018-98559], and NIGMS award number [R35GM133708] to I.M. The funders had no role in the design or execution of the study. The content is solely the responsibility of the authors and does not necessarily represent the official views of the National Institutes of Health. The UK Biobank Resource was used under Application 33923. The Health and Retirement Study genetic data were accessed through dbGaP accession phs000428.v2.p2. The Health and Retirement Study is sponsored by the National Institute on Aging (grant numbers U01AG009740, RC2AG036495, and RC4AG039029) and was conducted by the University of Michigan. The Jackson Heart Study (JHS) data were accessed through dbGaP accession phs000286.v6.p2. The study is supported and conducted in collaboration with Jackson State University (HHSN268201800013I), Tougaloo College (HHSN268201800014I), the Mississippi State Department of Health (HHSN268201800015I/HHSN26800001) and the University of Mississippi Medical Center (HHSN268201800010I, HHSN268201800011I and HHSN268201800012I) contracts from the National Heart, Lung, and Blood Institute (NHLBI) and the National Institute for Minority Health and Health Disparities (NIMHD). The authors also wish to thank the staffs and participants of the JHS. Funding for CARe genotyping was provided by NHLBI Contract N01-HC-65226. The Women’s Health Initiative (WHI) data were obtained from dbGaP accession phs000200.v12.p3. The WHI program is funded by the National Heart, Lung, and Blood Institute, National Institutes of Health, U.S. Department of Health and Human Services through contracts HHSN268201600018C, HHSN268201600001C, HHSN268201600002C, HHSN268201600003C, and HHSN268201600004C. This manuscript was not prepared in collaboration with investigators of the WHI, has not been reviewed and/or approved by the Women’s Health Initiative (WHI), and does not necessarily reflect the opinions of the WHI investigators or the NHLBI. Funding for WHI SHARe genotyping was provided by NHLBI Contract N02-HL-64278.

## Web Resources

Analysis code, https://github.com/mathilab/Height_Prediction_PRS

Neale Lab Summary statistics (GWAS round 2, accessed April 2, 2019), https://www.nealelab.is/uk-biobank) LiftOver, https://genome.ucsc.edu/cgi-bin/hgLiftOver

