## Supplementary Figures for "Polygenic scores for height in admixed populations"

### Supplementary Material

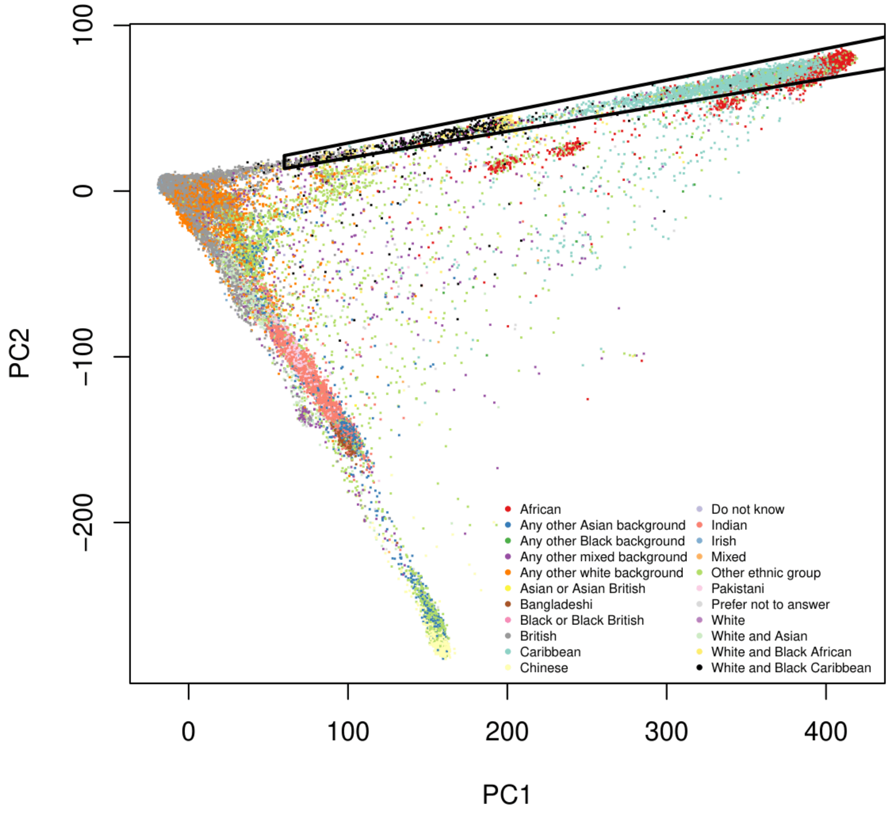


**Figure S1. Principal component analysis (PCA) of UK Biobank individuals.** PC1 separates Africans and Europeans. We selected 8,813 individuals with African or admixed African and European ancestry based on PCA, shown in the boxed area.

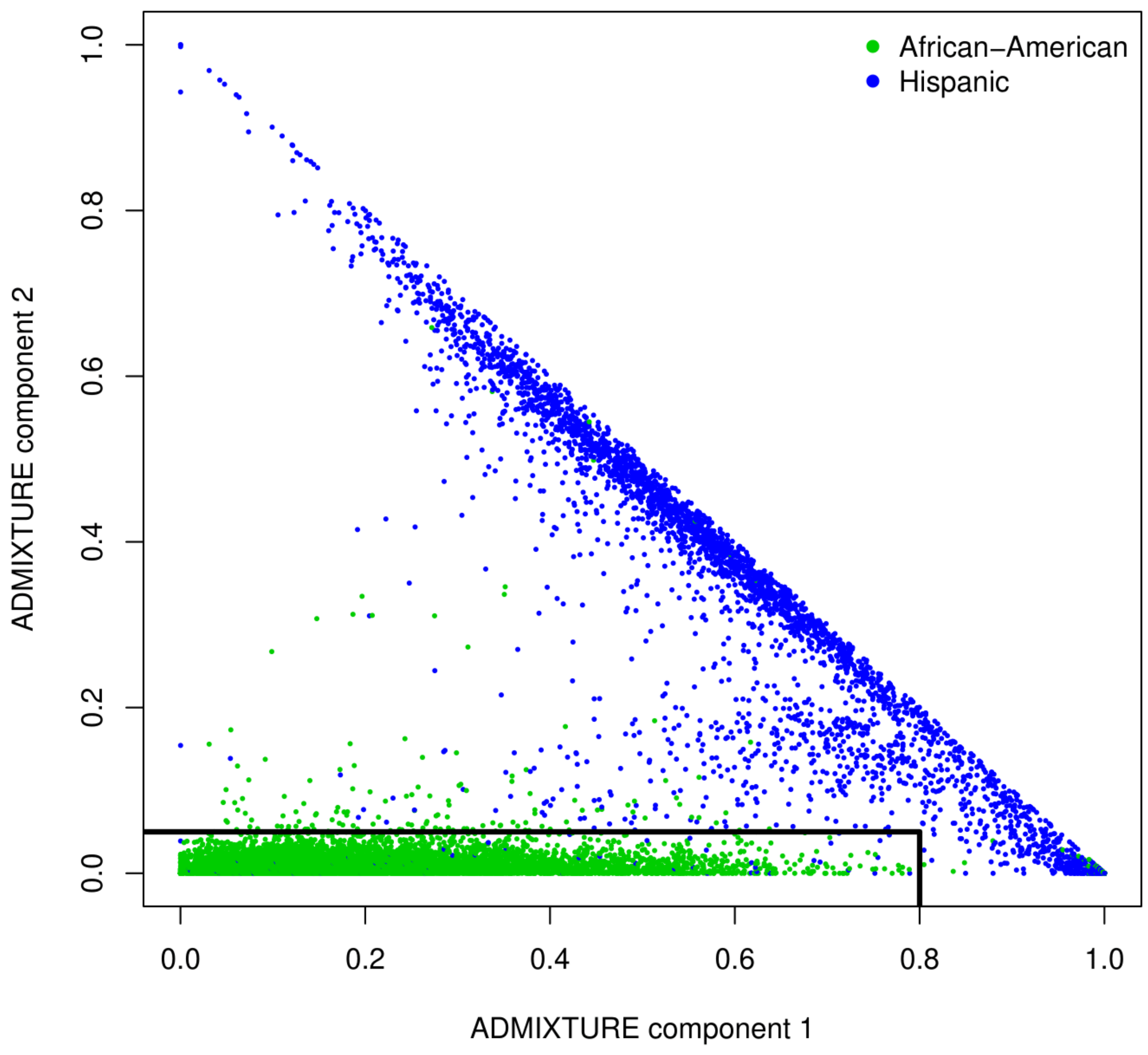


**Figure S2. ADMIXTURE analysis of Women's Health Initiative (WHI) individuals.** We ran unsupervised ADMIXTURE (k=3) and identified 7,285 individuals with self-reported "African American" ancestry with at most 0.8 of the first ADMIXTURE component (interpreted as the European component) and at most 0.05 of the second ADMIXTURE component (interpreted as the Native American component).

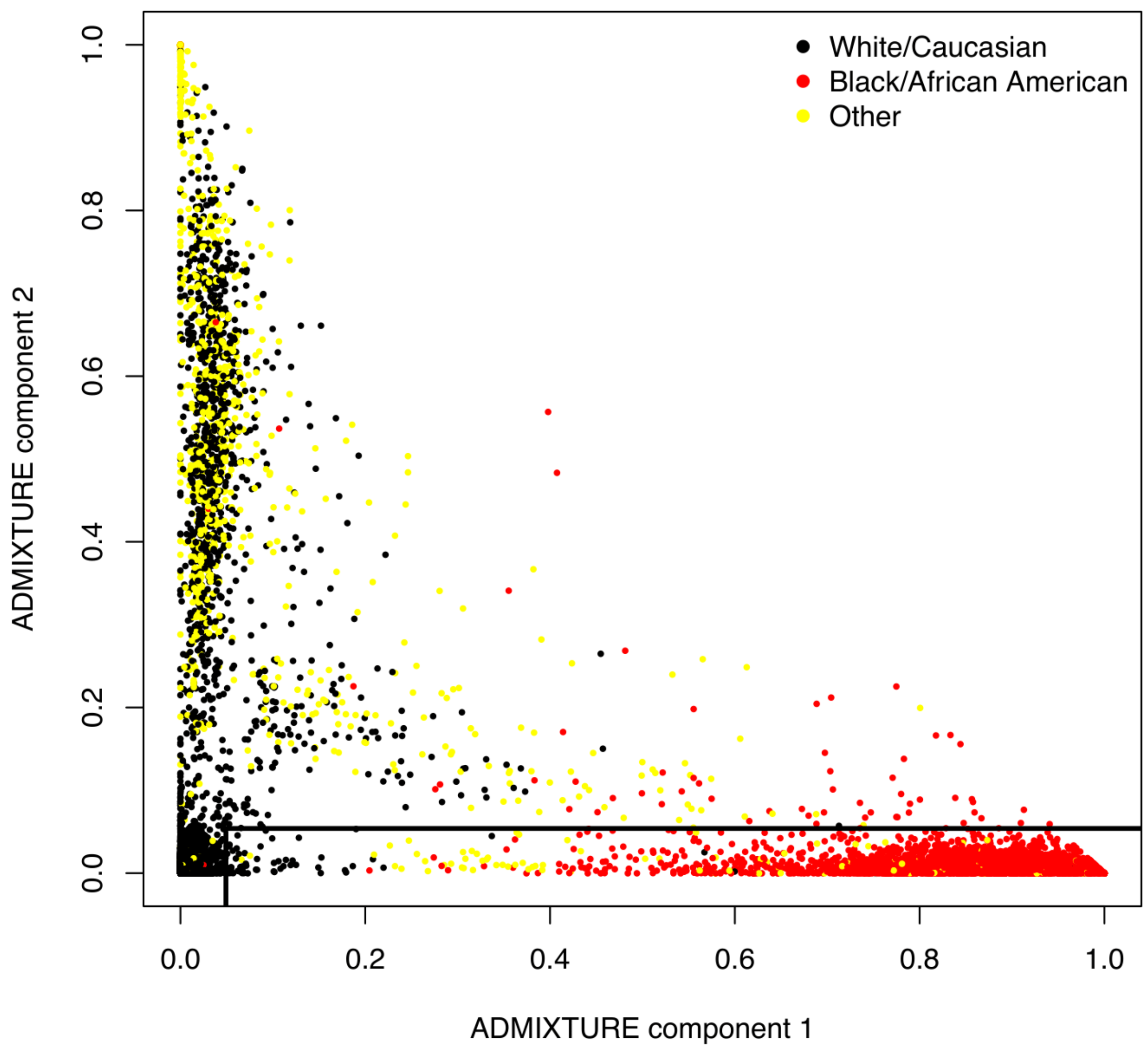


**Figure S3. ADMIXTURE analysis of Health and Retirement study (HRS) individuals.** We ran unsupervised ADMIXTURE (k=3) and identified 2,322 individuals with self-reported "Black/African American" ancestry with at least 0.05 of the first ADMIXTURE component (assumed to be African) and at most 0.05 of the second ADMIXTURE component (assumed to be Native American).

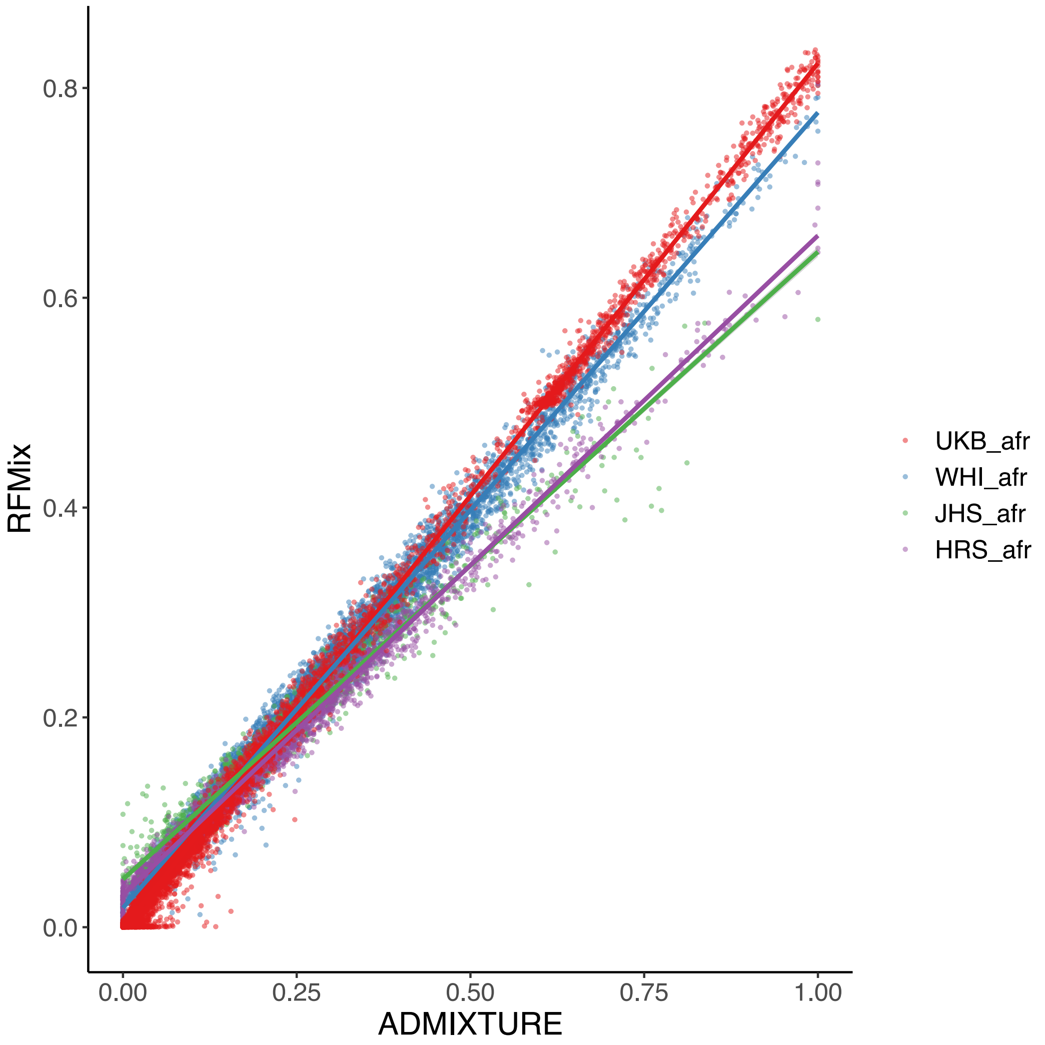


**Figure S4. Comparison between ADMIXTURE and RFMix estimates for genome-wide ancestry.** For RFMix, the average across all sites was used. Colored lines display linear regressions for each dataset. HRS_afr, Health and Retirement Study African Americans (*N*=2,322); WHI_afr, Women's Health Initiative African Americans (*N*=7,285); JHS_afr, Jackson Heart Study African Americans (*N*=1,774); UKB_afr, individuals with European and African admixture in the UK Biobank (*N*=8,813). Correlations are: 0.994 (WHI_afr), 0.982 (JHS_afr), 0.999 (UKB_afr), 0.993 (HRS_afr), 0.993 (ALL). The RFMix proportions are more likely to be correct because of the fact that we selected individuals with at least some African Ancestry, which will bias the ADMIXTURE results. Empirically, the RFMix proportions show a spike at 50% ancestry – likely to be from individuals with one parent of African ancestry and one of European ancestry. This spike is at ~60% European ancestry in the ADMIXTURE proportions suggesting that the ADMIXTURE European ancestry proportions are overestimated by a factor of around 1.2.

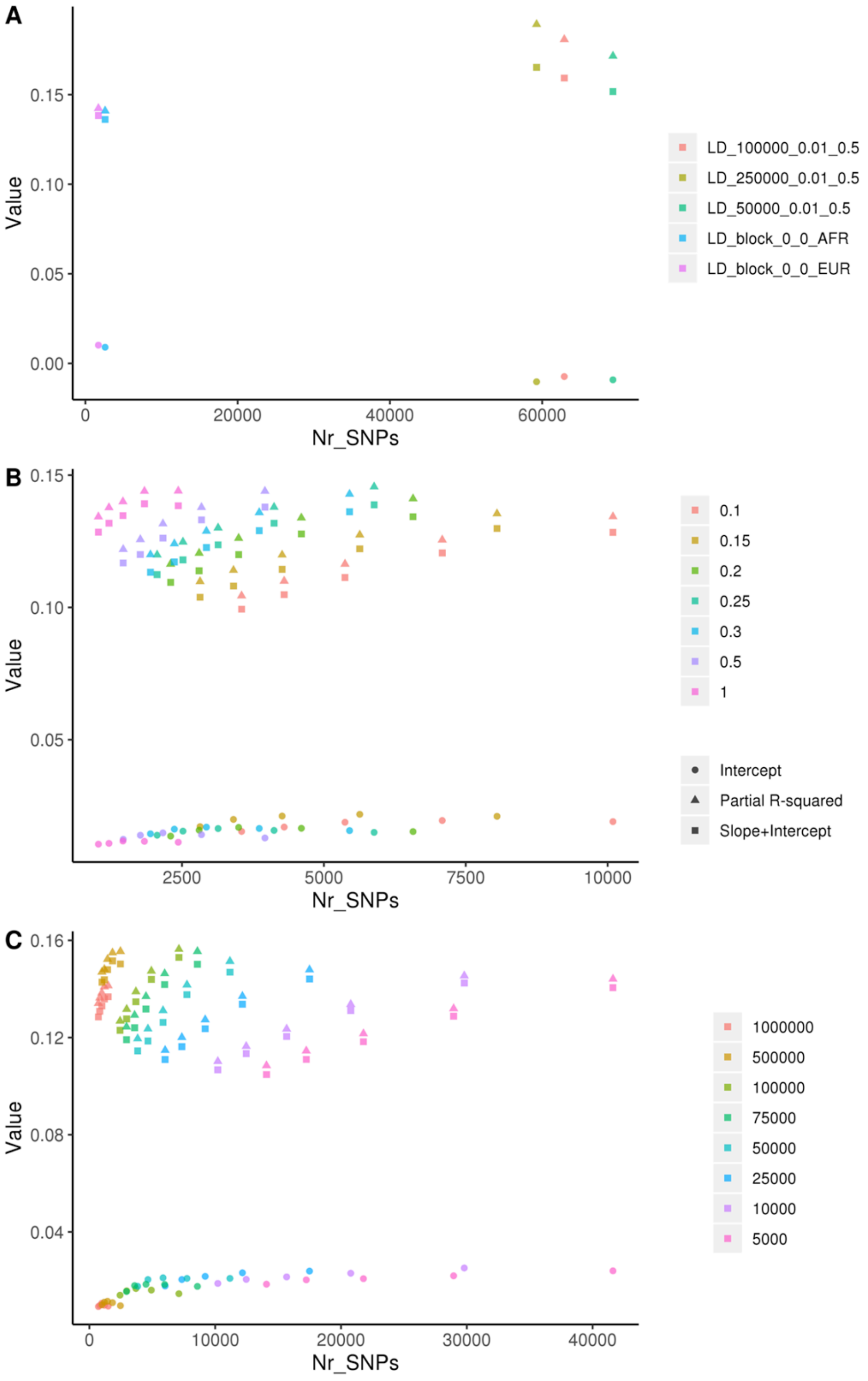


**Figure S5. Predictive power of height PRS as a function of number of SNPs included in the PRS. A**: LD clumping methods ($r^2$>0.5, p<0.01, window sizes 50-250 Kb) and LD blocks pruning strategy[3]. **B**: Genetic and **C**: physical clumping. Values are for the HRS_eur dataset.

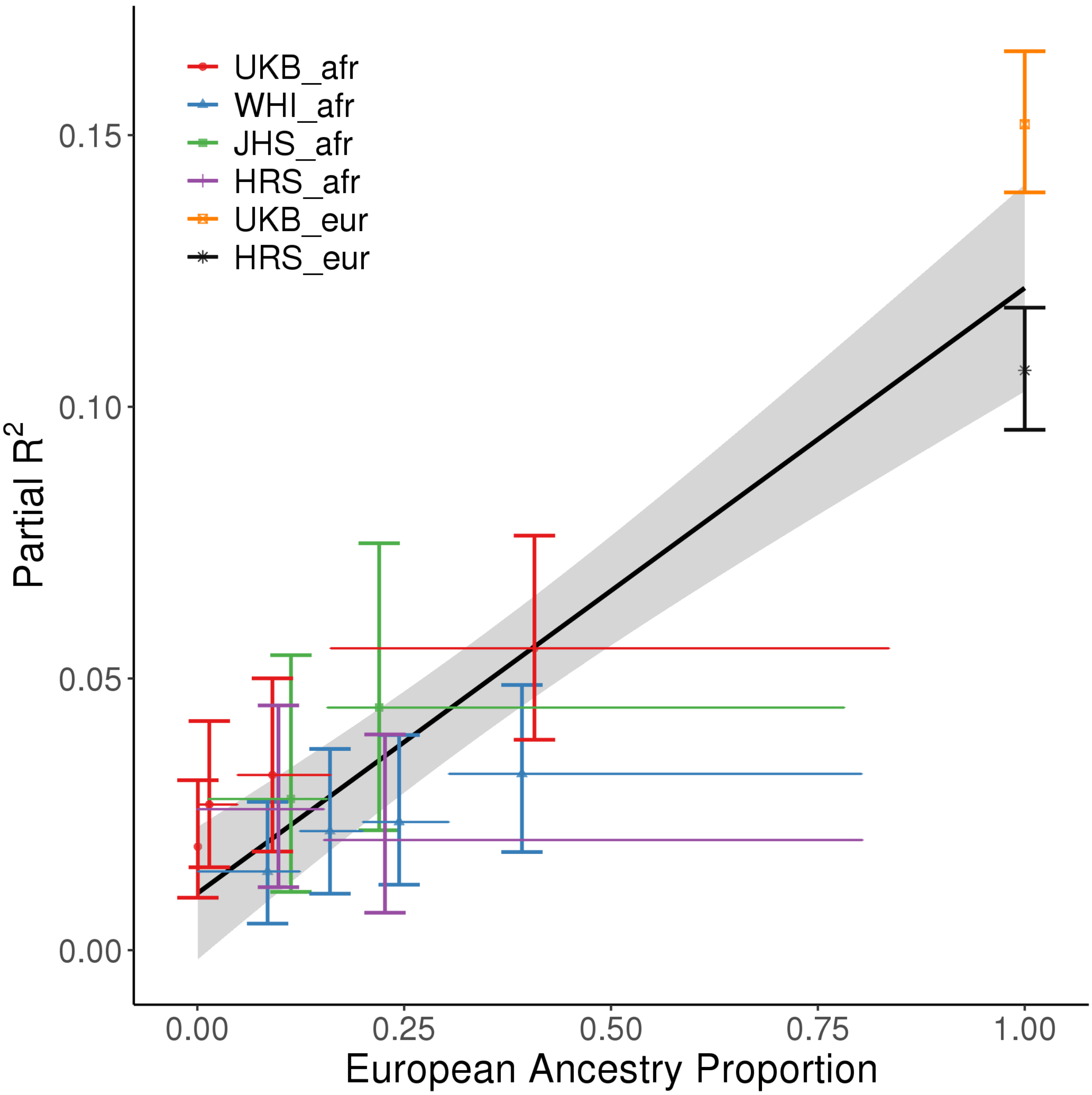


**Figure S6. As Figure 1, but using effect sizes re-estimated within sibling pairs.** Each admixed population is split up by quantiles of ancestry. Each cross represents a single quantile bin. Vertical bars represent bootstrap confidence intervals for partial-$R^2$. Horizontal bars represent the range of European ancestry included in that bin.

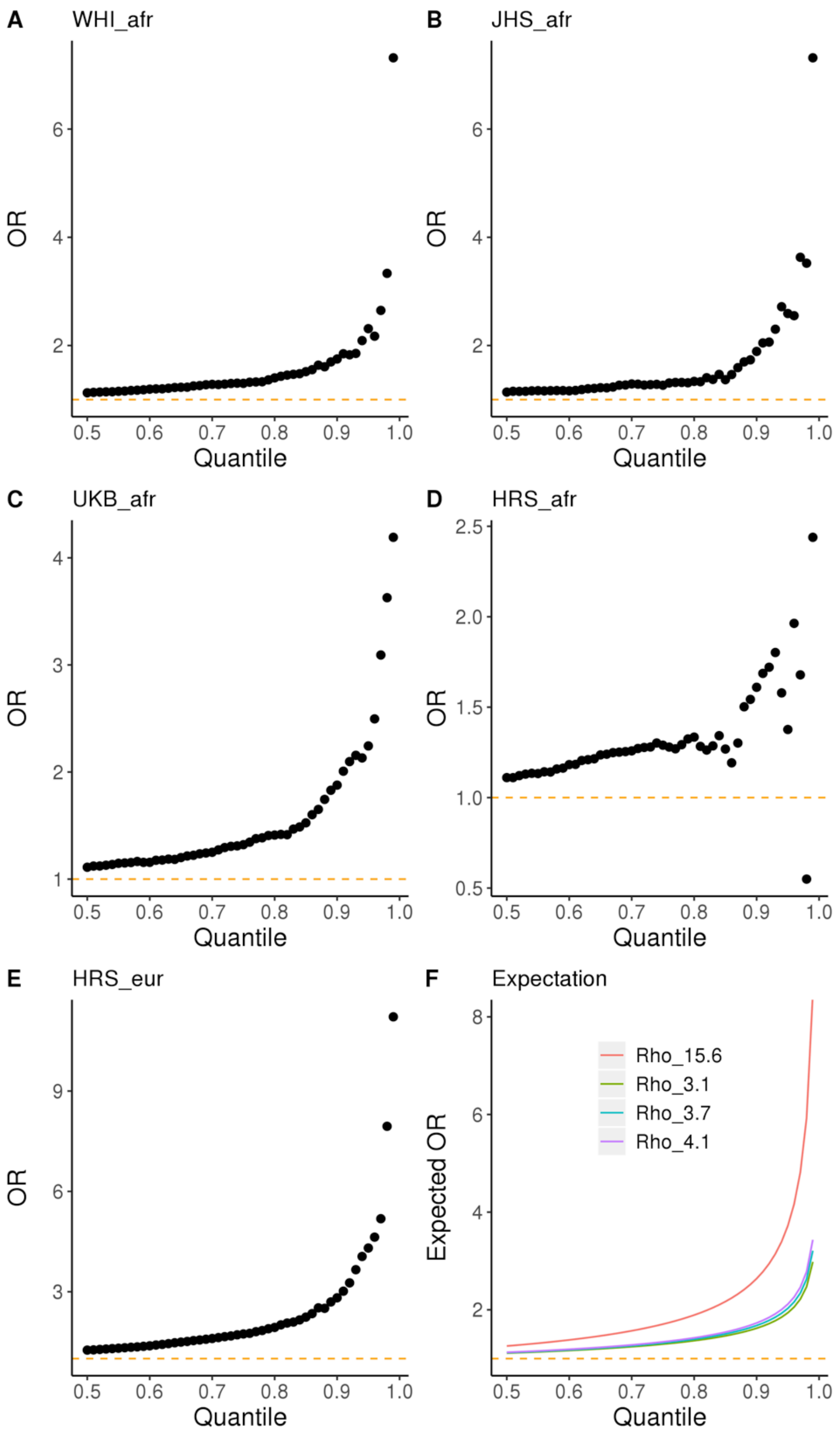


**Figure S7. Ability of PRS to predict "tallness". A-E**: Odds-ratio (OR) for being above the *q*-th quantile of height, conditional on being above the *q*-th quantile of PRS, for different datasets. **F**: expected OR assuming a bivariate normal distribution for PRS and disease risk on the liability scale. Different *rho* values reflect partial-$R^2$ detected empirically for each dataset (0.156 for HRS_eur, 0.041 for UKB_afr, 0.36 for WHI_afr, 0.38 for JHS_afr and 0.31 for HRS_afr).

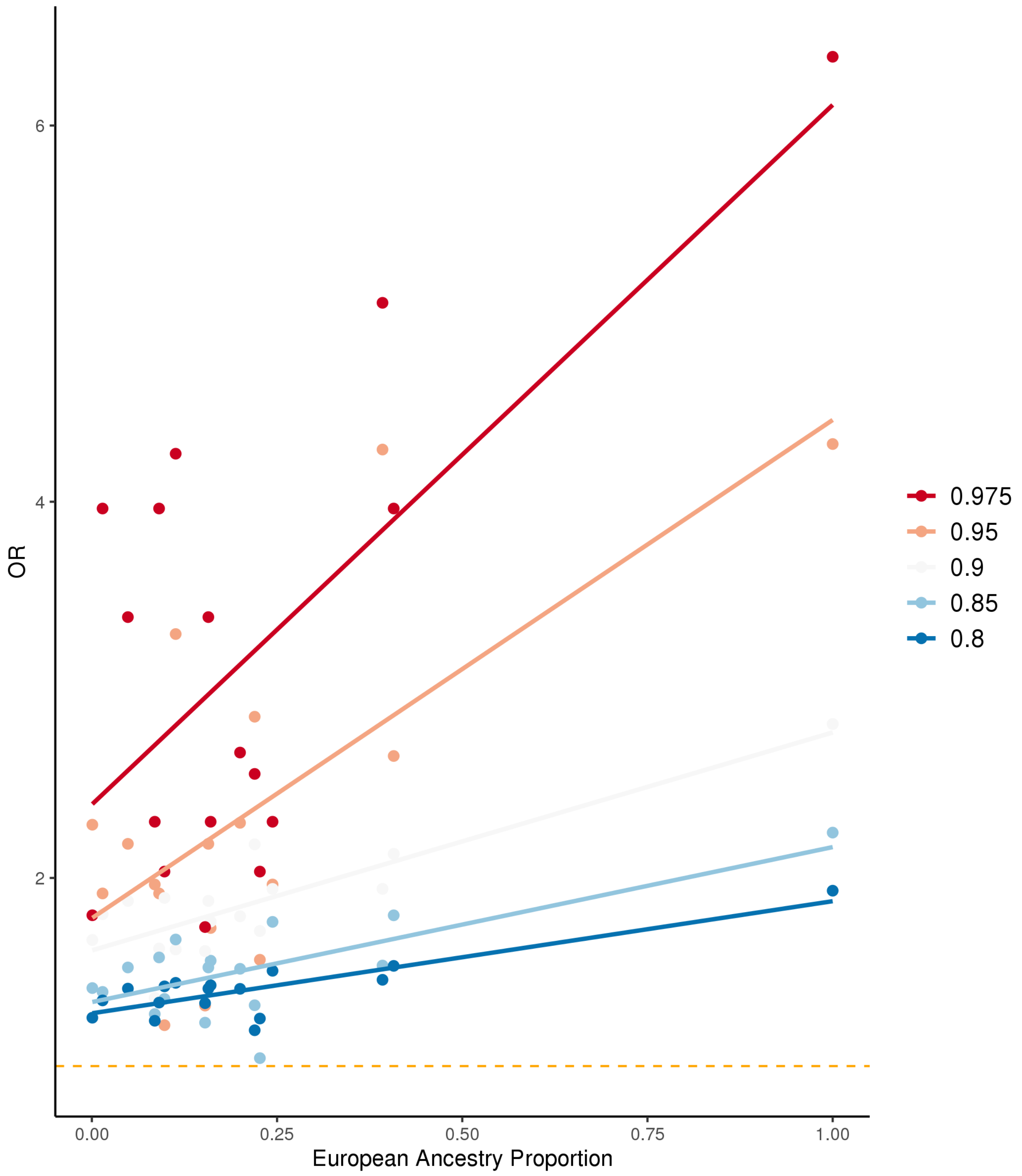


**Figure S8. Odds-ratio for 'tallness' as a function of European ancestry proportion.** All datasets are pooled together. OR is the probability of being in above the $n^{th}$ quantile of residual height (after regressing out sex, age, $age^2$, study) given that an individual is above the $n^{th}$ quantile of PRS, divided by 1-). Dashed line is OR=1.

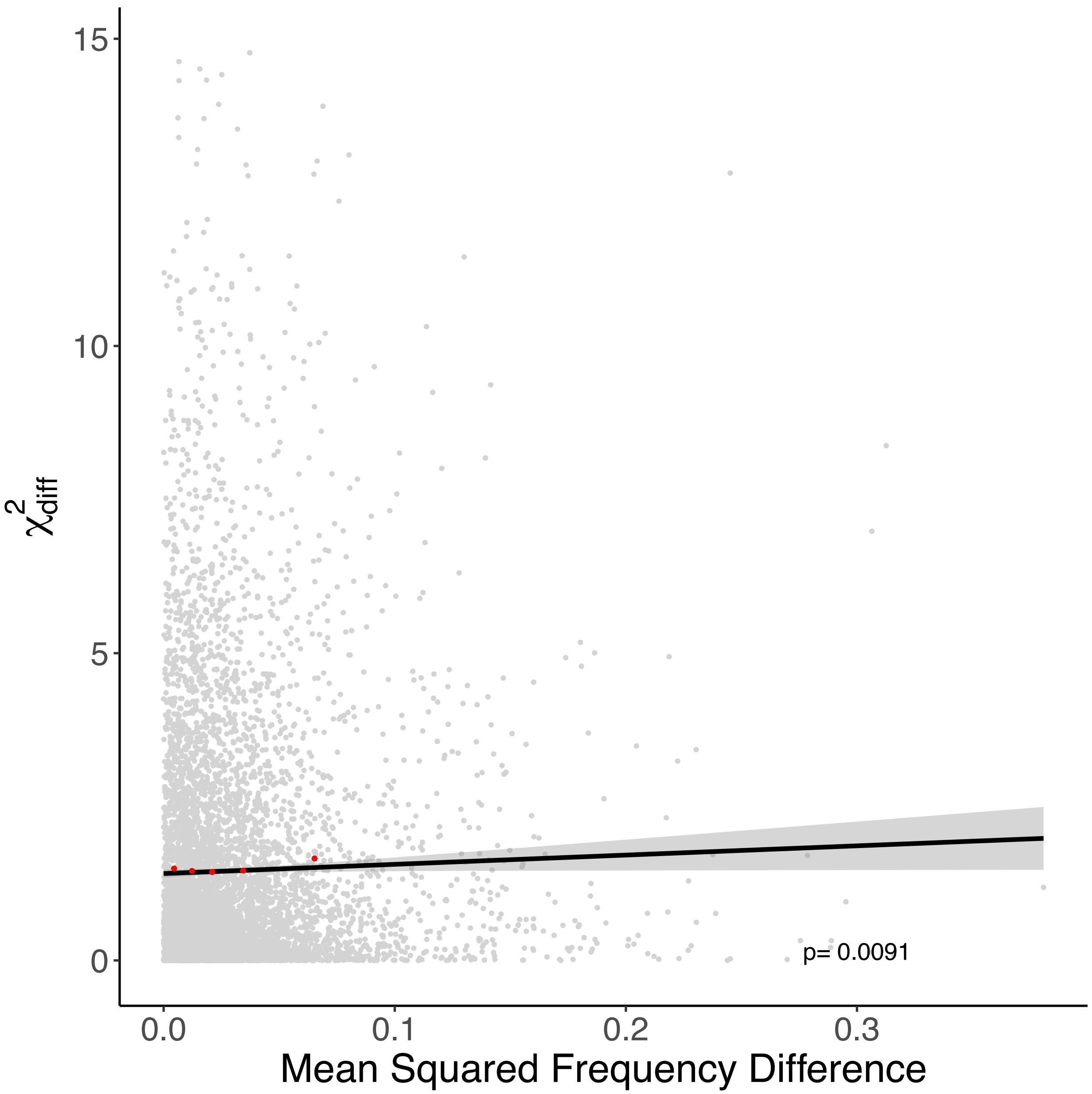


**Figure S9. Differences in effect size as a function of allele frequency difference around PRS SNPs.** x-axis, mean squared frequency difference for PRS SNPs for EUR and AFR in a 10 Kb window around each PRS SNP. Frequencies were calculated per dataset (HRS_eur, HRS_afr, UKB_eur, UKB_afr) for the causal allele. y-axis, $\chi^2_{diff}$ of the difference in betas estimated for EUR and AFR. Cut-off at 15 for display purposes excludes 15 data points. In red, for visualization, are median squared frequency difference values for 5 bins of mean squared difference and the mean $\chi^2_{diff}$ for each bin.

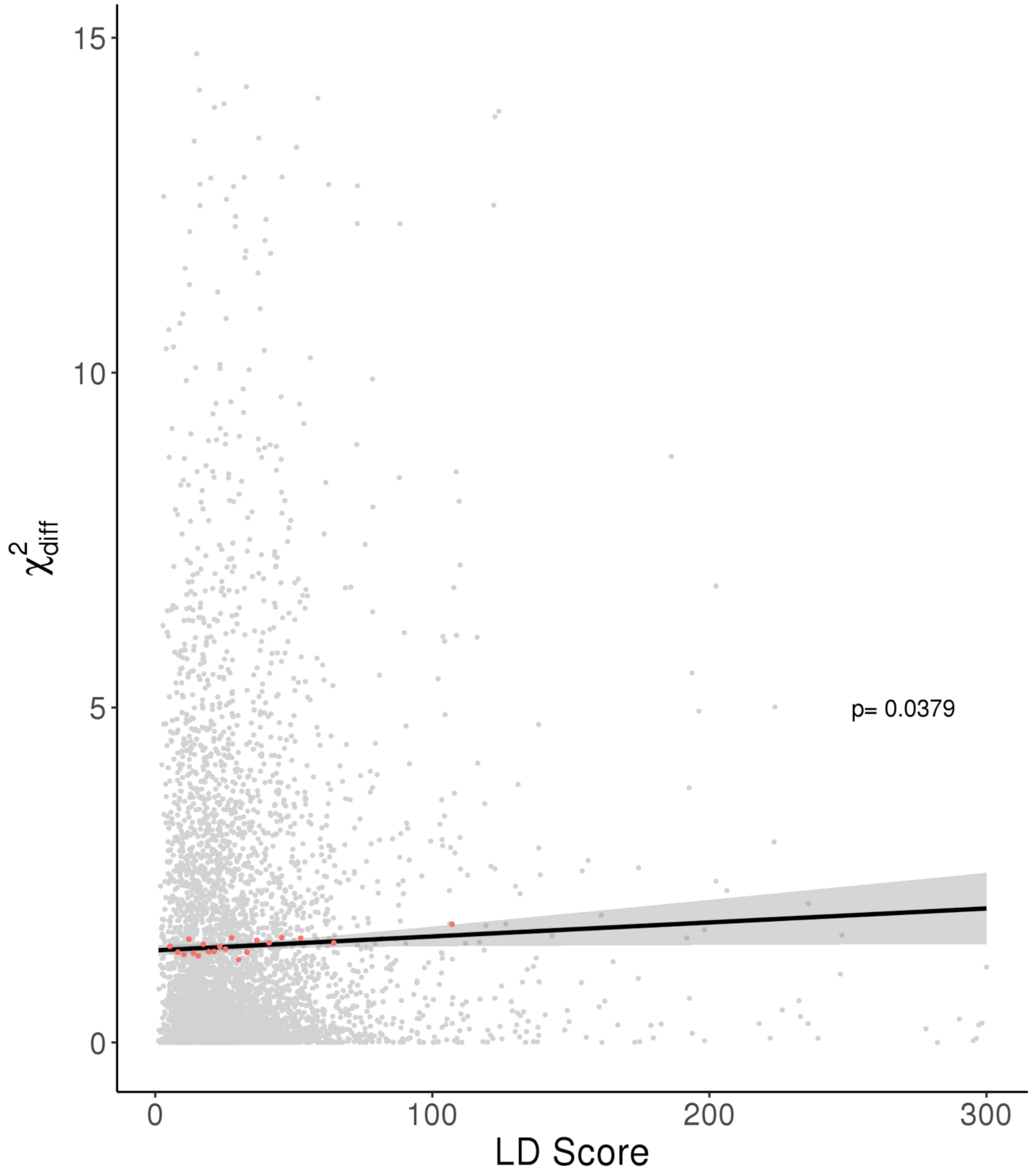


**Figure S10.** $\chi^2_{diff}$ against European LD score. Black line is the linear regression with confidence intervals in gray. Red points represent median recombination distance and LD score values for 20 bins and mean $\chi^2_{diff}$ values.

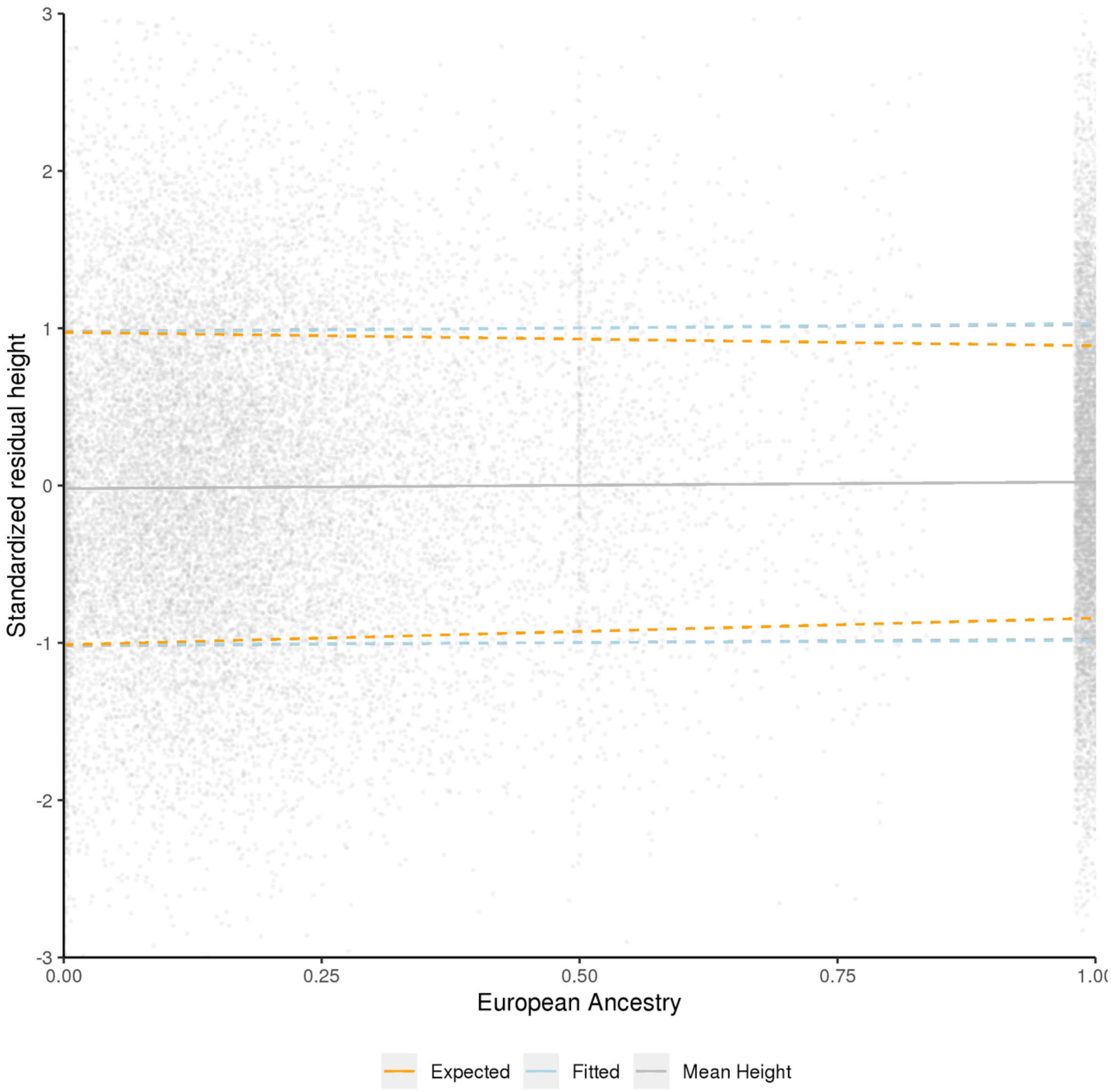


**S11. Phenotypic variance.** Gray lines show mean height and dashed lines show ∓1 sd. In orange we show the expected sd if it were negatively dependent on European ancestry. In blue we show the fitted model with variable variance, which is not significantly different from the constant variance model (green). We reject the model (orange line) where the phenotypic variance in people with 100% European ancestry is 76% that of people with 0% European ancestry.

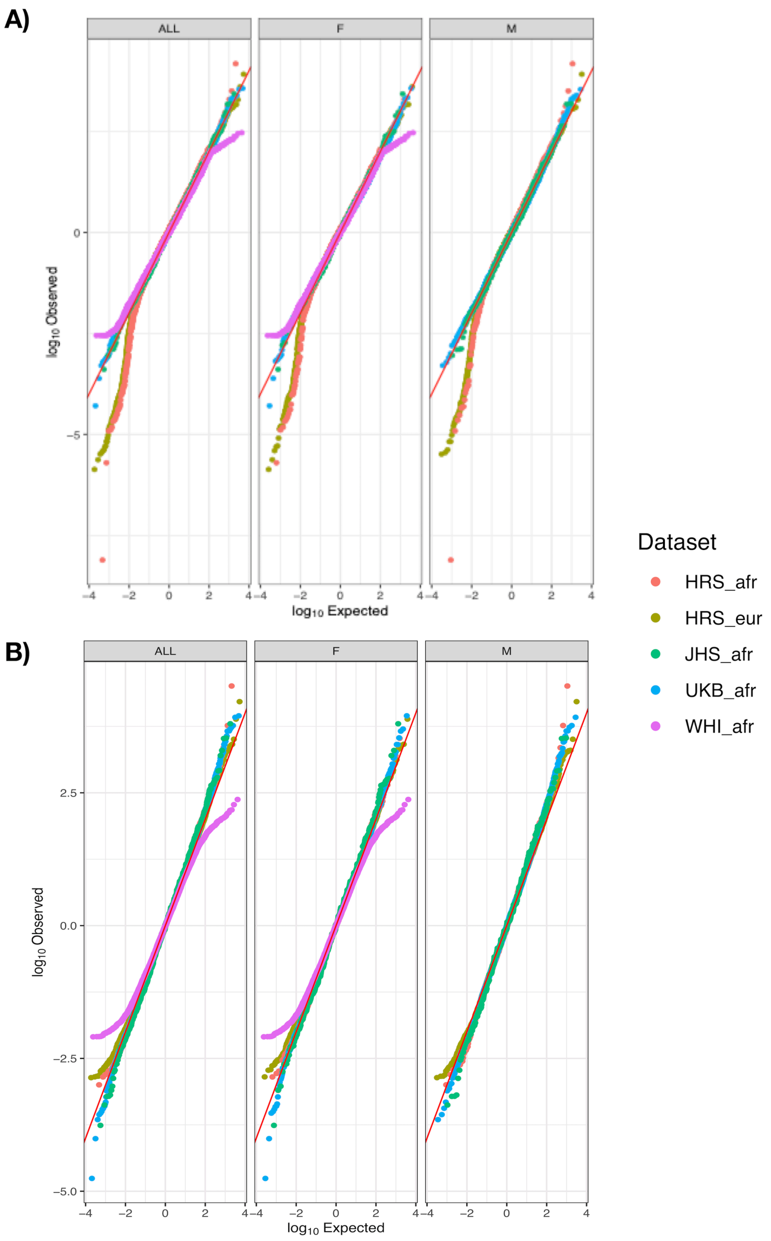


**Figure S12. Q-Q plot of residual height in each dataset**. **A:** before filtering; **B:** after filtering out individuals from WHI and HRS datasets according to the following criteria. WHI: individuals with height more than 2 standard deviations away from the mean; HRS_afr and HRS_eur: individuals with height lower than the mean minus 2 standard deviations (mean and sd calculated sex-specific). X-axis, expected values under a normal distribution. F, females; M, males; ALL, males and females. Residuals obtained after regressing out confounding factors as follows: height~sex+Dataset+sex*Dataset+age+age* Dataset+age*sex+age$^2$+age$^2$*Dataset+age$^2$*sex.
